# Comparing Robotic and Computer Vision Assessments of Unilateral and Bilateral Reaching in Healthy Adults

**DOI:** 10.1101/2025.08.05.668208

**Authors:** Desmond Asante, Shelby Ziccardi, Stephen Guy, Rachel L. Hawe

## Abstract

Assessment of reaching is foundational to upper limb neurorehabilitation. Current neurorehabilitation needs have increased the demand for quantitative clinical assessments of bilateral coordination. Robotics and computer vision for motion tracking are two means to provide relevant quantitative metrics but have many differences including the dimensionality of reaching movements (planar versus three-dimensional) and data acquisition. We do not know how consistent measures of bilateral coordination performance are between these different assessments. In this study, we examined how one robotic and one computer vision method can identify differences between symmetrical and asymmetrical reaching, and the correlations in movement time, and hand lag between these two approaches. Thirty healthy young adults completed four reaching games using the Kinarm exoskeleton robot and a custom developed augmented reality assessment using computer vision.

We found that both approaches were able to detect well-established movement time and hand lag differences between symmetrical and asymmetrical reaching, with the differences between symmetrical and asymmetrical being larger with the computer vision approach. Moderate correlations were found between approaches for unilateral and symmetric reaching in both movement time and hand lags; however, no significant correlations were found between approaches for asymmetric reaching.

Our results show that reaching task performance differs between robotic and computer vision-based assessment, however, both approaches provide quantitative metrics of unilateral and bilateral reaching that are consistent with prior research. There are benefits and tradeoffs to each approach, and this study informs how clinicians and researchers can consider the methodological differences when determining which assessment method to use.

## 1. Introduction

Reaching is a primary way we connect with our environment. Depending on the task, we reach unilaterally or bilaterally, with bilateral reaches requiring either symmetric or asymmetric coordination between the limbs (Kantak et al., 2017). Neurological injuries such as stroke impair both unilateral and bilateral reaching, leading to chronic limitations in performing daily activities (Park, 2023; Sainburg et al., 2013). Even after long term rehabilitation, up to 50% of stroke survivors are unable to gain a moderate level of upper limb function (Langhorne et al., 2011; Mayo et al., 1999; Sathian et al., 2011).

A barrier to improved rehabilitation outcomes is that clinicians lack quantitative measures for assessing reaching impairments after stroke. Traditional clinical measures rely primarily on subjective observations and focus on the more affected arm (Barreca et al., 2006; Barrett et al., 2013; Chen & Bode, 2010; Krumlinde-sundholm & Eliasson, 2003; Shirota et al., 2016). Due to the subjective nature of current clinical assessments, clinicians face a challenge in prescribing targeted treatments and tracking patient progress, ultimately delaying functional upper limb recovery. The Stroke Recovery and Rehabilitation Roundtable (SRRR) has emphasized the critical need for technology-based assessments and quantitative metrics to facilitate upper limb recovery post-stroke. As part of the upper limb assessments post-stroke, the SRRR highlighted the importance of quantifying 2D planar reaching movements using a robotic device and 3D functional tasks including reaching for and drinking from a cup (Kwakkel et al., 2017, 2019). Since stroke impairments have been shown in both the contralesional and ipsilesional arms (Meyer et al., 2016; Lai et al., 2019) as well as with bilateral coordination, a limitation of these approaches is not including a bilateral component. Assessment of bilateral reaching can detect impairments in both arms individually, as well as assess an individual’s ability to coordinate the two limbs together, which is not detected by unilateral assessments.

In alignment with SRRR’s recommendations, robotic systems have been employed to provide objective measures of reaching abilities. While many commercially available robotic systems are unilateral, the Kinarm exoskeleton robot has been used to quantify both unilateral and bilateral movements in healthy and neuroatypical populations (Poitras et al., 2024; Scott & Dukelow, 2011). Robotic systems constrain movements to the horizontal 2D plane, which has the benefits of simplifying biomechanical analyses and enabling movement in individuals with more severe motor impairments, however, the movements may differ from unconstrained and more functional movements. Additionally, most robotic systems are not readily available in the clinic due to cost and complexity. More recently, computer vision-based markerless motion tracking systems have also been explored (Colyer et al., 2018; Debnath et al., 2022; Huber et al., 2024; Friedrich et al., 2025). Though these systems overcome restrictions of marker-based motion tracking, they are still not widely used clinically. This is partly because they inherit similar hardware complexities as the marker-based 3D motion capture systems such as requiring multiple cameras and complex analysis pipelines that do not easily translate to routine clinical use. We have previously developed an integrated computer vision and augmented reality approach to assess reaching abilities requiring only a single camera (webcam) and computer (Ziccardi et al., 2024). This system requires minimal setup and measures reaching movements not constrained to a 2D plane, which may make it a more clinically feasible quantitative assessment compared to robotics. However, we do not know how reaching performance may differ when individuals perform planar reaches in a robotic device versus three dimensional reaches with this set up.

Beyond the type of reaching movements and acquisition methods, the choice of metrics impact clinical utility. Traditional clinical measures use ordinal scales, which raise concerns regarding their validity, susceptibility to floor effects and ceiling effects, and low inter-rater reliability (Balu, 2009; Carlsson et al., 2003; Duncan et al., 1997; Ghandehari, 2013; Lien et al., 2023; Peters et al., 2015; Stewart & Cramer, 2013). In contrast, quantitative assessment metrics have shown strong reliability and validity, while providing meaningful clinical insights (Bigoni et al., 2016; Latreche et al., 2023; Scott & Dukelow, 2011; Wagh et al., 2024). However, quantitative metrics are overabundant. A 2019 systematic review reported 151 different upper limb kinematic metrics (Schwarz et al., 2019). While an array of kinematic metrics can increase insights of motor control and identify specific impairments in clinical populations, clinicians have a need for simple metrics that do not require intensive computation, can be used to detect impaired overall performance, and be sensitive to change with treatment or recovery. Movement time can capture factors including speed, efficiency, and corrective movements, while hand lags (i.e. difference between individual movement times of both arms during bilateral reaching) provide meaningful measures of interlimb coordination in bilateral reaching (Rose & and Winstein, 2013; Schwarz et al., 2019).

To support the growing need for quantitative metrics, our lab has developed both robotic and computer vision-based assessment methods. These two approaches may be suitable for different settings, with robotics able to provide fine-grained detailed data in research laboratories while computer vision approaches are more likely to be utilized in the clinic. In this study, we compared temporal measures (movement times and hand lags) of unilateral and bilateral reaching tasks using the Kinarm exoskeleton robot and a computer vision/augmented reality system (adapted from Ziccardi et al., 2024). These systems are not intended to be direct substitutes, as they have differences in biomechanical requirements and perceptual attributes, but we expect both to be able to quantify unilateral and bilateral reaching and identify similar key differences. As a first step in establishing our computer vision approach as a valid assessment of reaching abilities, the objective of this study was to determine how the performance of healthy young adults compares between robotic and computer vision-based assessments of unilateral and bilateral reaching. Specifically, we examine how each method can identify differences between symmetrical and asymmetrical bilateral reaching, as there are well established neurophysiological and kinematic differences (Aramaki et al., 2010; Blinch et al., 2014, 2015). We also examine the correlations in movement time and hand lag between the two approaches. We hypothesized that each method could detect movement time and hand lag differences between symmetrical and asymmetrical reaching. We also hypothesized that for unilateral and symmetrical reaching, there would be at least a moderate correlation between robotic and computer vision-based assessments.

This study will determine the ability of both robotic and computer vision approaches to quantify unilateral and bilateral reaching, as well as demonstrate performance differences between the two approaches. As computer vision approaches are more clinically feasible than robotics, this will inform us of the pros and cons of such quantitative assessments.

## 2. Methods

### 2.1 Participants

Thirty participants were recruited and enrolled from the University of Minnesota community. The sample size was estimated based on our unpublished preliminary data using a similar approach (modified in this study) with both robotic and computer vision assessments. We had found an effect size of Cohen’s d = 0.9 for the robotic and d = 0.83 for the computer vision assessments. Based on these effect sizes, a significance level of α = 0.05 and a statistical power of 0.80, minimum sample sizes of 10 and 11 were estimated using G*Power version 3.1.9.7 (Faul et al., 2007). We increased this minimal sample to 30 participants due to multiple comparisons and correlation analyses.

Participants were included if they were between 18 and 35 years and had no history of any motor or sensory disorders limiting their upper limb movements. After screening, participants reported to the NeuRAL lab at the University of Minnesota where they were consented. Participants’ age, sex, and self-reported hand dominance, defined by writing hand, were collected. The average age of participants was 25.6 ± 4.9 years; 15 were males and 15 females; 29 were right dominant and 1 was left dominant.

### 2.2 Procedure Overview

Participants’ unilateral and bilateral reaching were tested with both the robotic and computer vision assessments, in a single session in a randomized order. Participants were allowed breaks in between the two assessments. The computer vision assessment session for each participant took approximately 10 minutes while the robotic assessment took approximately 30 minutes to complete including set-up. This research was reviewed and approved by the Institutional Review Board (IRB) at the University of Minnesota.

### 2.3 Robot assessment procedure

Participants sat in the Kinarm chair with their upper limbs supported in the horizontal plane by bilateral robotic exoskeletons to eliminate the effect of gravity, as shown in Figure 1A. The robot’s joints were aligned to participants’ shoulder and elbow joints, allowing for unrestricted movement in the horizontal plane. Participants viewed visual targets reflected from a horizontal screen to appear in the same plane as their limbs. Participants were blocked from seeing their actual arms by an opaque screen; however their fingertip position was represented virtually by a white dot at the tip of their index finger.

**Figure 1.**
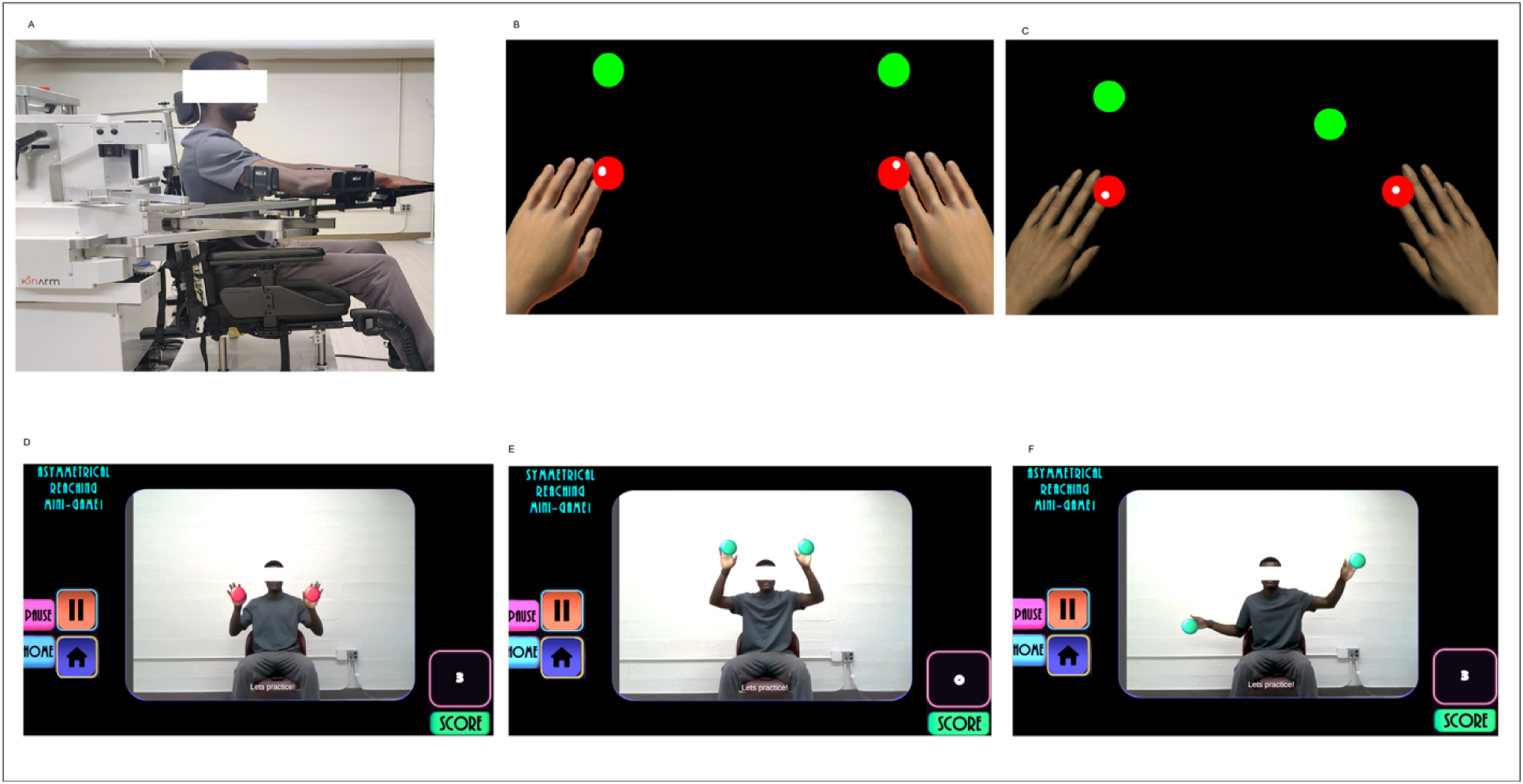
Robotic and computer vision methods. **A)** The participant is shown setup in the Kinarm exoskeleton with arms supported by bilateral exoskeletons in front of the horizontal display. **B and C)** The workspace for the robotic reaching tasks, where the red targets are start positions and green targets are end positions 12 cm away. The hand positions are shown by the white cursors. Participants do not see their arms while performing tasks. **B)** Task workspace for the symmetrical reaching. **C)** Task workspace for the asymmetrical reaching. **D-F)** The participant’s view on the game display while completing the computer vision tasks. Participants see themselves as well as two virtual targets displayed on the screen. Scores and the control buttons are the right and left side of the screen respectively. **D)** Start target position for bilateral computer vision assessment. **E)** Symmetric reaching end targets. **F)** Asymmetric reaching end targets. ***Photographs are of author of manuscript*.**

The robotic assessment involved four tasks in a randomized order, all requiring participants to move their arms either unilaterally or bilaterally to reach the targets shown in the display, as illustrated in Figure 1 B &C. All reaches began with participants positioning their fingertips in the red start targets with 2 cm radii, which required a starting position of 30° shoulder horizontal abduction and 90° elbow flexion. After a random time delay between 1800 and 4200 ms, peripheral targets 12 cm from the start target, turned on. The peripheral target coordinates were constant across all tasks, however used in different combinations. Unilateral dominant and non-dominant reaching required participants to reach for a single target at a time, with all targets displayed on the ipsilateral side (Figure 2A). Bilateral symmetric reaching involved reaching with both arms simultaneously to two targets located in mirrored positions (Figure 1B). Bilateral asymmetric reaching also required simultaneously reaching with both arms, however, the target positions differed by 45° (e.g. As shown in Figure 2A, when the left hand reached for target 2, the right reached for target 11). Participants then reached as quickly and accurately as possible to the end target(s). Participants held their hands at the peripheral targets for 1500ms before they turned off and the trial was complete. Prior to each task, the experimenter used both verbal instructions and video demonstrations to ensure that participants understood the task. Unilateral and symmetric tasks had 35 trials each (7 repetitions of each target location in a random order), while asymmetric tasks had 48 trials (6 repetitions of each target location in a random order). The greater number of trials for the asymmetric task was due to more combinations of target locations. Kinematic data was collected for offline analysis with a sampling rate of 1000 Hz.

**Figure 2.**
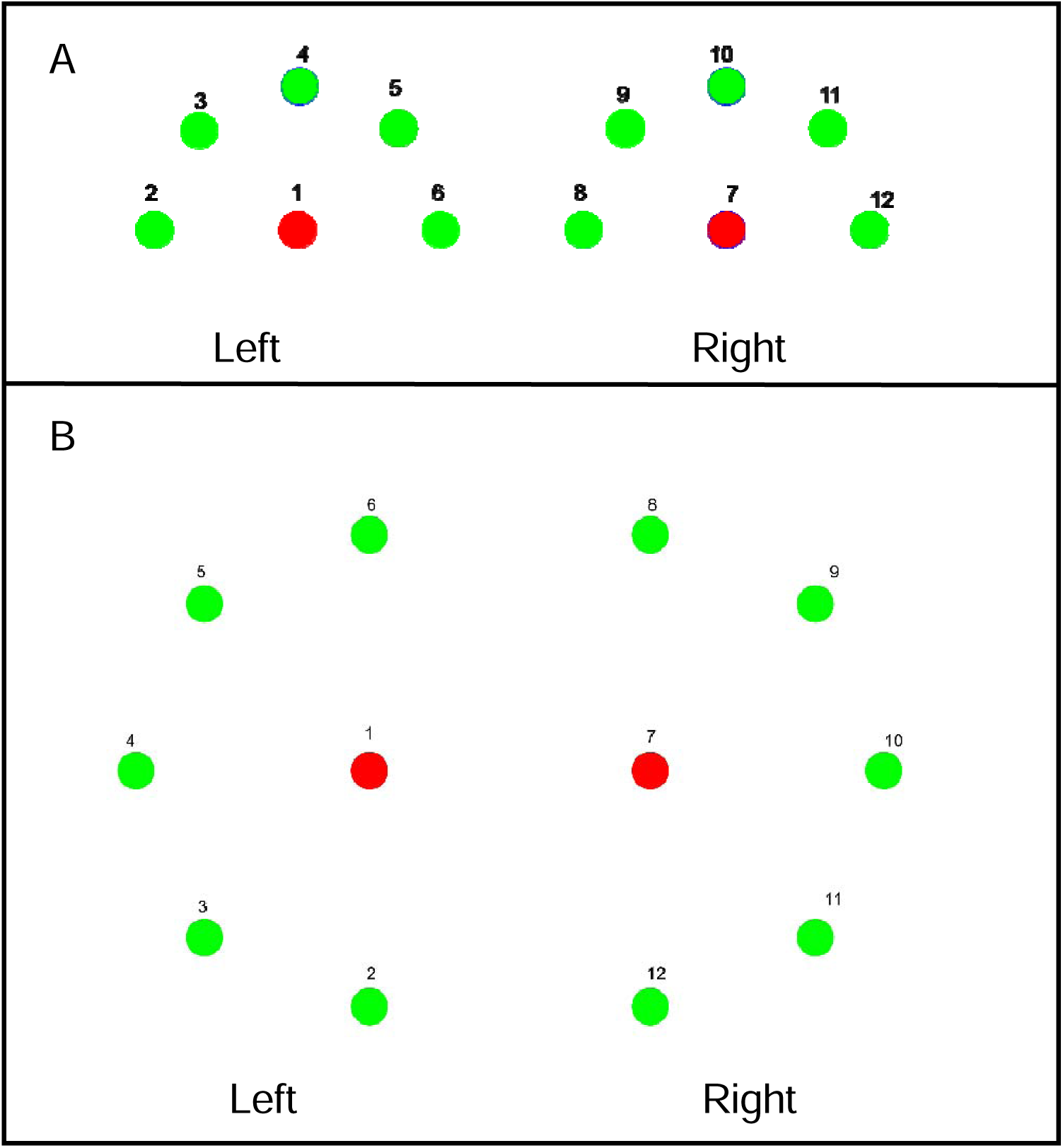
Target locations. Starting positions are shown in red with all possible end target locations shown in green. **A) Target positions for the robotic tasks as displayed on horizontal display.** Start and end targets are 12 cm apart. Unilateral tasks involved reaching with just the left hand (targets 1-6) or the right hand (targets 7 - 12). Symmetric tasks required both hands to reach mirrored targets (e.g.,1 & 7 to 4 & 10), while symmetric tasks involved reaching to different targets. (e.g.,1 & 7 to 3 & 10). All peripheral targets are approximately 12cm from the start target. **B) Target positions for computer vision tasks as displayed on vertical display.** Unilateral tasks involved reaching with either the left hand (targets 1–6) or right hand (targets 7– 12). Symmetric tasks required both hands to reach mirrored targets (e.g., 1 & 7 to 4 & 10), while asymmetric tasks involved reaching to different targets (e.g., 1 & 7 to 5 & 10). Distance between start and end targets was scaled to approximately 25% of participant’s wingspan. The distance between start and end targets was an arbitrary length of 0.5 within the normalized range.

### 2.4 Computer vision assessment procedure

In this assessment, participants moved their arms unilaterally and bilaterally to reach virtual targets while their hand position was tracked using MediaPipe for human pose estimation (Latreche et al., 2023; Wagh et al., 2024). The task was developed using the Unity 3D game development engine (version 2021.3.16f1). The task was an adaptation of a task previously used by Ziccardi et al, with adaptations made to standardize starting positions, require the same number of reaches per participant rather than as many as possible in a set time, and added in unilateral reaching (Ziccardi et al., 2024).

Initially, participants sat on an armless, stationary chair, backdropped by a plain gray wall. They faced a computer monitor (*32 in, 2560 ×1440 resolution*) with a webcam (*720p Logitech C720*) mounted on top that was used to track participants’ movements. Participants could see both their reflection as well as virtual targets on the monitor (Figure 1B). Participants were centrally positioned such that the center of the midline of the sternum corresponded to the midline of the display screen. To calibrate reach distance to approximate wingspan, the participant was moved forward and backwards with their chair until their full wingspan (shoulders abducted at 90 degrees) horizontally spanned the gameplay area displayed on the monitor. The webcam was then adjusted so that their full overhead reach (shoulders flexed at 180 degrees) reached the top edge of the gameplay area displayed on the monitor.

The assessment involved four tasks in a randomized order to match the robotic protocol-unilateral dominant and non-dominant reaching, bilateral symmetric reaching, and bilateral asymmetric reaching (Figure 2B). For each task, participants were instructed to move as fast and accurately as possible, reaching from start targets (red circles) to peripheral targets (green circles). Each time a participants’ hand in their onscreen reflection reached the start target as it was displayed on the monitor, it disappeared and was replaced by a peripheral target, and vice versa. The start targets were displayed at participants’ shoulder level. The peripheral targets were displayed in a random order at five different locations, equidistant to the start targets as shown in Figure 2B. Each target location was repeated 6 times in a random order for 30 total trials for each of the four assessments. To minimize the confounding effect of fatigue, participants took a break after fifteen trials for approximately one minute and resumed when they were ready. Before each task, participants performed five reaching trials to become familiar with the augmented reality system and ensure understanding of the task. Kinematic data was sampled at approximately 60 frames per second.

### 2.5 Data Analysis

Robotic kinematic data was low-pass filtered using a third-order, double-pass Butterworth filter implemented with cutoff frequency of 10Hz. Computer vision kinematic data was sampled from MediaPipe pose estimation and metrics were computed at runtime.

In both robotic and computer vision assessments, our primary outcomes were movement time for all tasks and hand lags for the bilateral tasks. These metrics were selected as they can be derived from events within the assessment task, and do not require analyzing kinematic data which may be both unreliable with a single camera and lack clinical feasibility due to increased computational demands. We defined movement time as the time the end targets appeared to the time the end targets were reached by the participant’s hand(s). For bilateral tasks, movement time ended when the second hand reached the target. Giving priority to clinical feasibility, we did not separate reaction time from the movement time as this would require analyzing kinematic data to determine the movement onset. In the symmetric and asymmetric tasks, absolute and relative (signed) hand lags (defined as the difference in reaching time of the dominant and non-dominant hand) were used as measures of bilateral temporal synchrony. Absolute hand lag was used to assess how synchronized the limbs were regardless of which hand reached first. Relative hand lag, which maintained the sign, was used to determine bias towards one hand leading. A positive relative hand lag means the non-dominant hand reached the target first. For each participant, median movement times and hand lags across trials were determined. Medians were used because the trial-level data was not normally distributed.

Outliers defined as values greater than 3 standard deviations away from the mean were removed. One outlier each was found in asymmetric movement times and symmetric and asymmetric hand lags. We used the Shapiro-Wilk Test to analyze the normality of variables (p > 0.05)(Shapiro & Wilk, 1965). As it is well established that asymmetrical movements have longer movement times and greater asynchrony between the hands, we determined if both methods can detect differences in the two types of bilateral reaches. We used paired t-tests and Cohen’s D, to determine differences and effect sizes between symmetric and asymmetric reaching for each assessment method. We then converted the movement times and hand lags from both assessments into z-scores. Since the two assessments are not designed to be identical, we used z-scores to preserve each measure’s relative position in its distribution for meaningful comparison. Movement time and hand lags for robotic assessment metrics were correlated with movement time and hand lags of computer vision assessment metrics using Pearson correlation. Lastly, as we expected that there may be a perceptual difference based on how far apart the targets are (robotic targets appearing further apart than computer vision targets due to participants’ distance from the display screen), we tested within the robot the association between movement time and distance between end targets using the linear mixed effect model. All data were processed and analyzed using MATLAB R2024a (Mathworks Inc., Natick, MA).

## 3. Results

### 3.1 Movement time

For the robotic assessment, the mean movement time for unilateral dominant reaching was 823 ± 173ms, while non-dominant reaching was 813 ± 159ms. Mean movement times for bilateral tasks are shown in Figure 3A. For bilateral symmetric reaching (Figure 3A), mean movement time was 1063 ± 262ms, while bilateral asymmetric (Figure 3A) was 1114 ± 222ms. Robotic assessments showed a significant difference between symmetric and asymmetric movement times (p = 8.1 x 10^-3^) with an effect size of d = 0.520.

**Figure 3.**
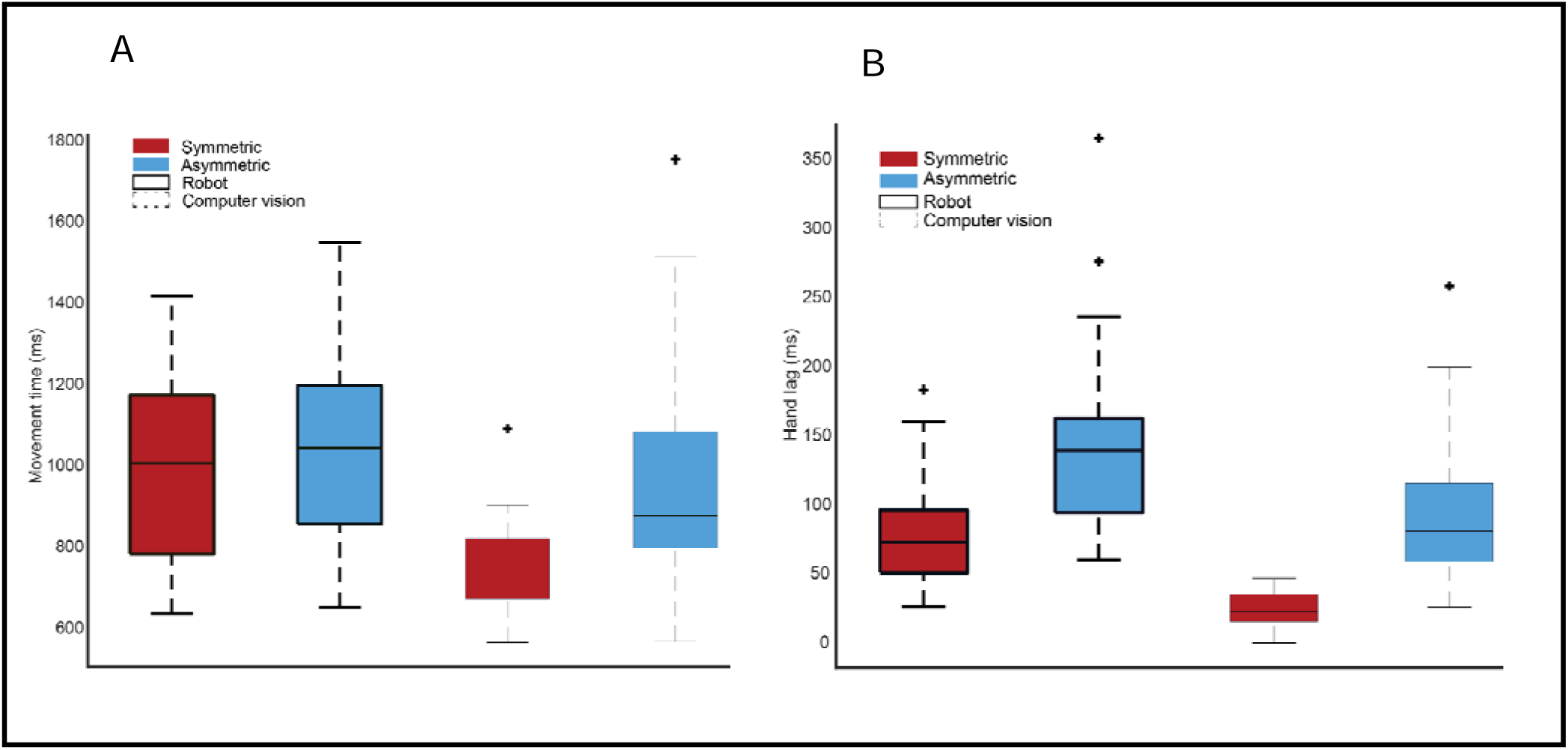
Comparison of movement time and absolute hand lag within robotic and computer vision methods. A) comparisons of movement time. B) comparisons of absolute hand lag. The thick-lined boxes represent robotic assessment, while the thin-lined boxes represent computer vision assessment. The red boxes represent symmetric reaching while the blue boxes represent asymmetric.

For the computer vision assessment, the mean movement time for unilateral dominant hand reaching was 698 ± 111ms, while non-dominant hand reaching was 710 ± 90ms. For bilateral symmetric reaching (Figure 3A), mean movement time was 822 ± 136ms, while bilateral asymmetric (Figure 3A) was 1103 ± 372ms. There is a significant difference between symmetric and asymmetric movement times (p = 3.9 x 10^-6^) with an effect size of d = 1.030.

Figure 4 shows the correlation between the z-scores of movement time between the robotic and computer vision assessments. In both the dominant and non-dominant reaching tasks (Figure 4), there was a significant moderate positive correlation between the movement times measured by the robotic and computer vision assessments (dominant: r = 0.450, p = 0.012, non-dominant: r = 0.400, p = 0.030). For the bilateral symmetric reaching task (Figure 4C), there was also a significant moderate positive correlation between the movement times measured by the robotic and computer vision assessments (r = 0.494, p = 0.006). However, for the bilateral asymmetric reaching task (Figure 4D), the correlation between the movement times measured by the robotic and computer vision assessments did not reach significance (r = 0.352, p = 0.060).

**Figure 4.**
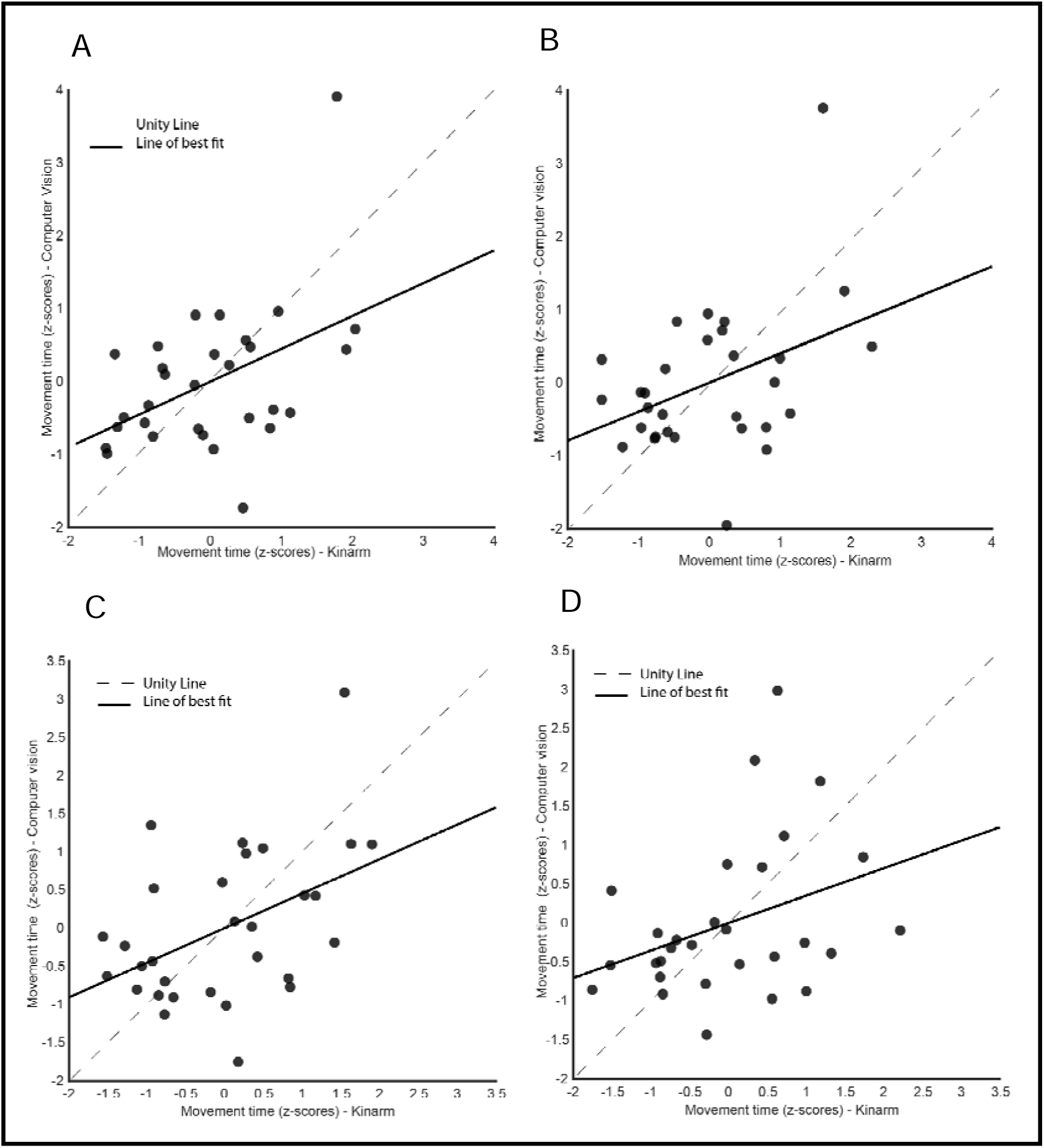
Correlation between robotic and computer vision assessment of movement time. In all four plots, the dashed lines represent the unity line (equal performance), while the solid line represents the line of best fit. The plots show movement times of unilateral dominant **(A)** and non-dominant hand reaching **(B)** as well as bilateral symmetric **(C)** and asymmetric reaching **(D)**.

### 3.2 Hand lag

With data from the robotic assessment, the mean absolute hand lag for bilateral symmetric reaching was 79 ± 38ms, while asymmetric (Figure 3B) was 142 ± 66ms. The mean relative hand lag for symmetric reaching was -5 ± 66ms, indicating the non-dominant hand reached first, while asymmetric was 6 ± 65ms, indicating the dominant hand reached first. Data from the robotic assessments showed a significant difference between symmetric and asymmetric absolute hand lags (p = 4.51 x 10^-4^) with an effect of size, d = 0.722.

With data from the computer vision assessment, the mean absolute hand lag for bilateral symmetric reaching was 22 ± 14ms, while asymmetric (Figure 3B) was 93 ± 51ms. The mean relative hand lag for symmetric reaching was 1 ± 9ms, while asymmetric was 13 ± 35ms, both indicating the non-dominant hand reached first. A significant difference between symmetric and asymmetric hand lag on the computer vision assessments was found (p = 1.210 x 10^-9^) with an effect size of d = 1.600.

Figure 5 A and B show the correlation between absolute hand lags in z-scores between the robotic and computer vision assessment. For bilateral symmetric reaching (Figure 5A), there was a significant moderate positive correlation between hand lags measured by the robotic and computer vision assessments (r = 0.566, p = 0.001). In contrast, for bilateral asymmetric reaching (Figure 5B), correlation between hand lags measured by the robotic and computer vision assessments did not reach significance (r = 0.060, p = 0.383).

**Figure 5.**
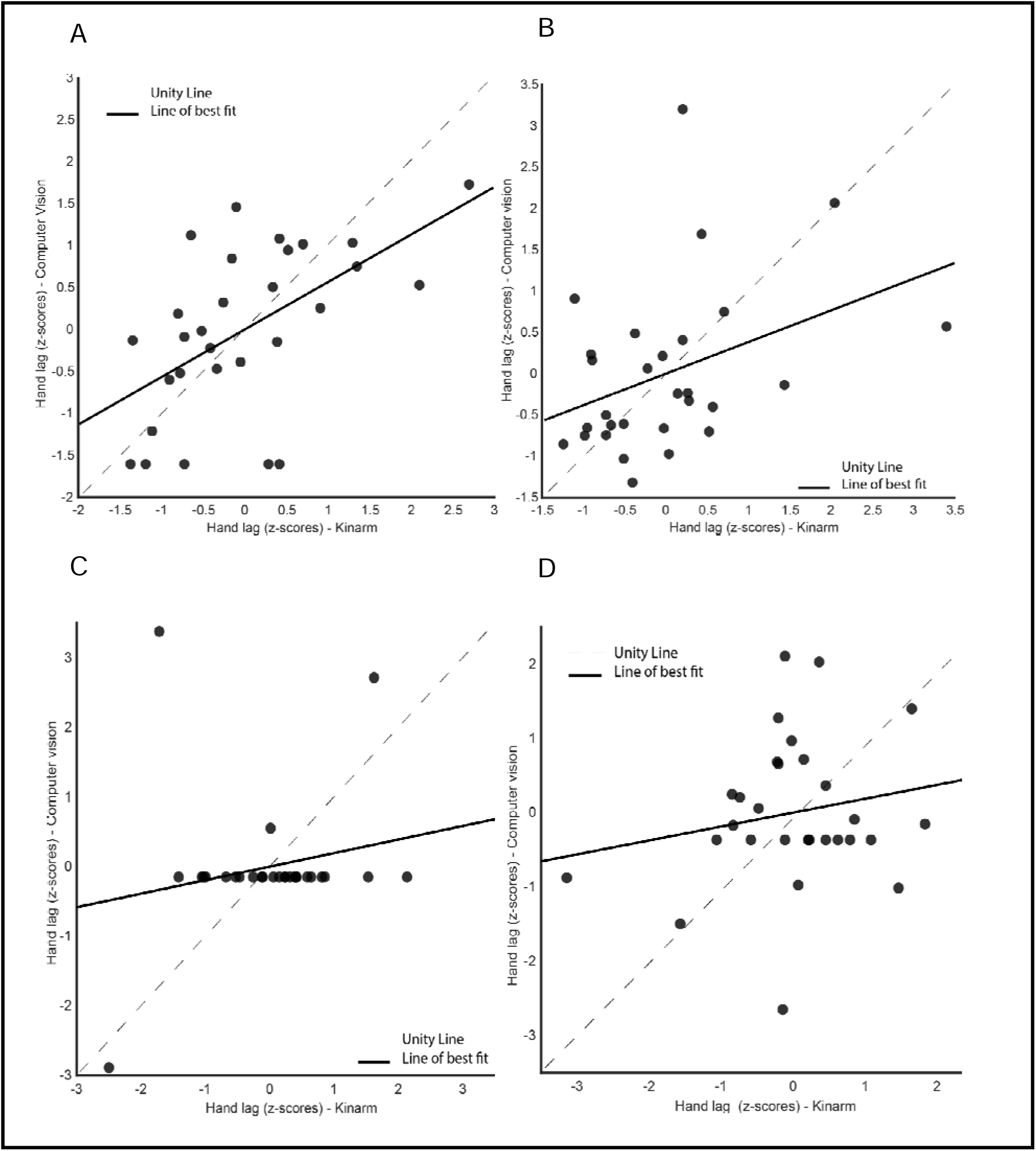
Correlation between robotic and computer vision assessment of hand lag. In all four plots, the dashed lines represent the unity line, while the thick line represents the line of best fit. The plots above show absolute hand lags for symmetric **(A)** and asymmetric reaching **(B)**. The plots underneath show relative hand lags of bilateral symmetric **(C)** and asymmetric reaching **(D).**

Figure 5 C and D show the correlation between the two methods for relative hand lags in z-scores. For both bilateral symmetric and asymmetric reaching (Figure 5), there was no significant correlation between hand lags measured by the robotic and computer vision assessments (symmetric: r = 0.195, p = 0.312, asymmetric: r = 0.187, p = 0.331).

### 3.3 Association between movement time and distance between end targets

There was a significant association between distance between peripheral targets and movement time in both symmetric (β= 3.373, p = 1.883 x 10^-6^) and asymmetric reaching tasks (β = 6.877, p = 1.459 x 10^-21^) as measured in the robotic assessment.

## 4 Discussion

In this study, we compared the performance of healthy young adults in robotic and computer vision assessments of unilateral and bilateral reaching. Examining temporal parameters, we found that both methods were able to identify previously well-established differences between bilateral symmetric and asymmetric movement times and hand lags. We found moderate correlations in movement times between robotic and computer vision assessments in unilateral and bilateral symmetric reaching tasks, however no significant correlation between methods for bilateral asymmetric reaching. These results demonstrate that while both approaches can provide quantitative measures of unilateral and bilateral reaching, much of how an individual performs is specific to the way reaching was assessed.

It is well established in both neurotypical populations across the lifespan as well as neurologically impaired populations that symmetric bilateral movements are the preferred pattern, which is reflected with faster movement times and more synchronicity or decreased hand lag (Blinch et al., 2014; Calabro & Perez, 2016; Gosser & and Rice, 2015). Therefore, our first approach in evaluating the two methods was to determine how they are able to differentiate symmetric and asymmetric reaching movements as a test of validity. In both methods, movement times and hand lags were shorter in symmetric than asymmetric reaching, giving validity to both approaches. The effect size for the difference between symmetrical and asymmetrical reaching was larger (d=1.03) for the computer vision compared to the robotic task (d=0.52). This suggests that our computer vision protocol was more sensitive to these different modes of motor control, for reasons later discussed.

Across all tasks, correlations between robotic and computer vision assessments were moderate at best, and not significant for bilateral asymmetric reaching. It is important to not interpret this result as one method being less valid than the other but rather reflects the specificity of each method. Although the tasks required symmetric and asymmetric arm movements, variations in the plane of movement, perceptual differences, and participants’ motivational affinity towards each method likely contributed to differences in reaching performance. Below we will discuss the different factors that influence how individuals may perform on each approach.

A primary difference in the approaches is the plane of movement. In the robotic assessments, participants moved horizontally in 2D, with arms supported, compensating for gravity (Poitras et al., 2024; Scott & Dukelow, 2011). The benefits of such a setup is that it simplifies biomechanics of reaching, and by providing arm support, reduces the confounding effect of fatigue and abnormal limb synergies that may impact neurologic populations such as stroke (Scott & Dukelow, 2011). However, the planar movements of the robotic device are often criticized for not matching the demands of functional real-world tasks. The computer vision approach can attempt to address this criticism by assessing reaching movements in 3 dimensions without limb support. By the nature of our approach and what is readily available with a traditional computer monitor, reaches with the computer vision system were displayed as vertical rather than horizontal reaches, thus having different biomechanical demands, although participants were not constrained to only vertical movement. The fact that we find differences between the two assessment strategies suggests that how people perform may be specific to limb support and movement plane. This is consistent with previous studies showing that gravity compensation and plane of movement alter upper limb motor control and kinematics in both healthy and clinical populations during reaching movements (Coscia et al., 2014; Krabben et al., 2012; Prange, Jannink, et al., 2009; Prange, Kallenberg, et al., 2009; Schaffer & Sainburg, 2017). It is therefore important in interpreting findings on unilateral and bilateral reaching as specific to the task that was performed, and measures may not generalize to an overall assessment on reaching abilities.

Beyond differences in the plane of movement, there were perceptual differences between the computer vision-based and the robotic assessment. The two systems differed in reaching amplitudes and target sizes due to what is feasible with each approach, with the robot having smaller target sizes. According to Fitt’s law, movement time is dependent on the both amplitude of movement and size of targets, and this law extends to bilateral reaching when sizes of two targets being reached are the same (J. A. S. Kelso et al., 1979; Fitts, 1954; Sardar et al., 2023). Additionally, visual distance between targets appeared shorter on the monitor used in computer vision assessment compared to the Kinarm display, due to the size of the monitor and distance from the screen, which we found to influence reaching performance. Another key difference is how participants viewed their arms. In the robotic task, participants’ fingertip positions were represented with dots, and they could not see their full limbs, while in the computer vision assessment participants could see their whole body on screen including their arms. Such differences in visual feedback have been shown to impact joint kinematics during reaching movements (Thomas et al., 2016). Our study confirms the fact that how people reach is affected by how they perceive targets.

Perceptual differences may explain why the differences between symmetrical and asymmetrical were greater for the computer vision approach, as well as why the two approaches were not correlated for the asymmetric tasks. The constraint due to the smaller target size in the robotic task likely alters the behavior and creates what has been described as a “hover” phase in which the two limbs get close to the targets but then hover while visual attention is switched between limbs to enable accuracy in the final contact (Miller & Smyth, 2012; Riek et al., 2003; Srinivasan & Martin, 2010). This causes the movement times to be slower in the symmetrical task for the robot approach, or closer to the movement times of the asymmetric task. Additionally, in the computer vision approach, while the accuracy constraint is lower, the targets are also closer together, allowing both to be viewed at the same time, decreasing the delay between the hands. When individuals do not have to be switching visual attention between hands to ensure accuracy in the computer vision assessment, the differences may better reflect differences in motor control between symmetric and asymmetric movements. However, this should not be interpreted as if one approach is superior or inferior to the other, as real life bilateral tasks vary in the distance between objects as well as accuracy constraints. Rather, it shows how the specificity of the task will impact performance.

Lastly, differences in participants’ intrinsic motivation and attention could have affected motor performance (Wulf & Lewthwaite, 2016). Subjective reports from participants suggest that they enjoyed the computer vision approach due to its more gamified nature. Though we randomized the assessment order and provided verbal encouragement during both, we believe that participants may have had differing motivational affinity towards each assessment, considering the methodological differences in affordances and constraints.

Despite differences in environmental constraints, each method determined notable differences between symmetric and asymmetric reaching metrics. This shows that overall, performance depends on the specifics of how bilateral coordination is assessed. Even using similar outcomes (movement time, hand lag) with the same nature of the task (unilateral and bilateral reaching), performance outcomes varied across methods. With the current advancements in technology assessment methods (Colyer et al., 2018; Debnath et al., 2022; Shirota et al., 2016), this finding challenges clinicians and rehabilitation scientists to determine standardized approaches for more consistency in how unilateral and bilateral reaching is assessed, as well as systematically investigate factors that may influence reaching assessments.

In this study, we limited our sample to healthy young adults to first determine how the two assessments differ for typical movement. Our next step is to use the two approaches in individuals with stroke and other neurologic conditions. Moving into clinical populations introduces additional benefits and tradeoffs for the two approaches. For individuals with stroke, the robot can support arms, which allows for the assessment of unilateral and bilateral reaching and coordination even with significant muscle weakness, and minimizes the abnormal synergistic arm movement commonly seen post-stroke. Individuals with moderate to severe stroke may have challenges performing the anti-gravity movements required by the computer vision approach due to weakness and flexion synergies, creating a floor effect. The location of targets may need to be individually scaled within an individual’s abilities; however, this causes challenges in comparing across participants.

As with other clinical assessments, both robotic and computer vision methods integrated virtual environments (Cóias et al., 2022; Faria et al., 2023). As confirmed in our study, performance outcomes in virtual environments are highly dependent on perceptual abilities(Höhler et al., 2021). To develop clinically feasible assessments, the perceptual limitations associated with stroke must be prioritized. In general, this calls for the design of assessment tools which can be personalized to the level of impairment of individuals with stroke. Again, towards clinical feasibility, the ease of this assessment must be a strong factor. In this study, we only focused on the temporal component of coordination. Temporal measures are easy to quantify and have been shown to be robust in investigating interlimb coordination (Rose & and Winstein, 2013). However, spatial measures like path lengths and hand trajectories also give a wealth of information on bilateral coordination (Tomita et al., 2017). Future work will use multiple cameras to derive spatial information for a more complete kinematic picture.

In conclusion, technology offers a way to assess bilateral coordination in a quantitative manner. Our findings show that reaching task performance differs between robotic and computer vision-based assessment. The robotic system is highly accurate but limits naturalistic arm movements and is not readily accessible clinically. In contrast, the computer vision-based assessment is easily accessible, and supports naturalistic movements but is susceptible to tracking inaccuracies due to camera set up. The trade-offs highlight the need for standardized technology-based methods that can be broadly applied to accelerate rehabilitation research and clinical assessments.

## Acknowledgements

This work was supported by the University of Minnesota McKnight Land Assistant Professorship to RLH and University of Minnesota Data Science Initiative Seed grant to SJG and RLH.

## CRediT authorship contribution statement

Desmond Asante: Writing – review & editing, Writing – original draft, Visualization, Software, Methodology, Investigation, Formal analysis, Conceptualization.

Shelby Ziccardi: Software, Methodology, Writing – review & editing.

Stephen J. Guy: Software, Methodology, Writing – review & editing, Funding acquisition, Supervision

Rachel L. Hawe: Writing – review & editing, Writing – original draft, Supervision, Methodology, Investigation, Funding acquisition, Conceptualization.

## Research Data

De-identified data is available in the data repository Zenodo. Due to participant’s faces being visible, raw video data is not made publicly available, but raw robotic data and analyzed data for the computer vision approach are shared on the repository. https://zenodo.org/records/16655664

